# Dynamics of repurposing synthetic and natural compounds against WNT/β-Catenin signaling in glioma- An *in vivo* approach

**DOI:** 10.1101/2022.12.02.518940

**Authors:** Daisy Precilla S, Shreyas S Kuduvalli, Indrani Biswas, Bhavani K, Agiesh Kumar B, Jisha Mary Thomas, Anitha T. S

**Affiliations:** Mahatma Gandhi Medical Advanced Research Institute (MGMARI), Sri Balaji Vidyapeeth (Deemed to-be University), Puducherry- 607 403, India; Department of Pathology, Mahatma Gandhi Medical College and Research Institute (MGMCRI), Sri Balaji Vidyapeeth (Deemed to-be University), Puducherry- 607 403, India; Catalysis and Energy Laboratory, Department of Chemistry, Pondicherry University, Puducherry- 605 014, India; Department of Biochemistry and Molecular Biology, Pondicherry University, Puducherry- 605 014, India

**Keywords:** Glioblastoma, Drug repurposing, WNT/β-catenin signalling, Apoptosis, Temozolomide

## Abstract

**Background:** Glioblastomas arise from multistep tumorigenesis of the glial cells and are associated with poor prognosis. Despite the current state-of-art treatment, tumor recurrence is inevitable. Thus, there exists a desperate need for effective therapeutic alternatives to improve glioblastoma outcome. Among the innovations blooming up, drug repurposing could provide a profound premises for glioblastoma treatment enhancement. While considering this strategy, the efficacy of the repurposed drugs as monotherapies were not up to par; hence, the focus has now shifted to investigate the multi-drug combinations to target glioblastomas. In line with this concept, we investigated the efficacy of a quadruple-combinatorial treatment comprising temozolomide (the benchmark drug) along with chloroquine (a synthetic drug), naringenin (a flavonoid) and phloroglucinol (a marine derivative) in an orthotopic glioma-induced xenograft model.

**Methods:** Anti-proliferative effect of the drugs was assessed by immunostaining. The expression profiles of WNT/β-catenin and apoptotic markers were evaluated by qRT-PCR, immunoblotting and ELISA. Patterns of mitochondrial depolarization was determined by flow cytometry. TUNEL assay was performed to affirm apoptosis induction. *In vivo* drug detection study was carried out by ESI-Q-TOF MS analysis.

**Results:** The quadruple-drug treatment had significantly hampered GB proliferation and had induced apoptosis by modulating the WNT/β-catenin signalling. Flow cytometric analysis revealed that the induction of apoptosis was associated with mitochondrial depolarization. Further the quadruple-drug cocktail, had breached the blood brain barrier and was detected in the brain tissue and plasma samples from various experimental groups.

**Conclusion:** The quadruple-drug combination served as a promising adjuvant therapy to combat glioma lethality *in vivo* and can be probed for translation from bench to bedside.

## 1. INTRODUCTION

Glioblastoma (GB) is recognized as one of the most lethal and highly infiltrative intracranial tumours, accounting for 29% of the central nervous system (CNS) malignancies (Lah et al. 2020). The current tripartite therapy for GB comprises maximal surgical resection followed by radiotherapy and administration of the DNA-alkylating agent, temozolomide (TMZ) (Davis 2016). Despite vast strides in the standard-of-care treatment, the median 5-year overall survival rate of GB patients remains to be less than 5% (Mangiola et al. 2010). Moreover, clinical data had demonstrated that GB patients require higher doses of TMZ, which in turn poses adverse toxic side-effects (Hart et al. 2013). Hence, there exists an unmet need of adjuvant therapies to alleviate TMZ-induced toxicity and concomitantly uplifting the sensitivity of GB cells to improve the therapeutic efficacy of the standard drug.

While the search for alternate therapeutic strategies to alleviate glioma pathogenesis is underway, previous studies had suggested that the overall prognosis in glioma patients can be improved by repurposing lead compounds of synthetic and natural origin, which in turn, would improve the therapeutic efficacy of TMZ (Xiao et al. 2016; Ashfaque et al. 2021; Precilla et al. 2022). These therapeutic regimes, as adjuvants to the standard drug, can hamper glioma progression via modulation of signalling pathways with tolerable side effects (Kim et al. 2010; Lu et al. 2012; Wang et al. 2016; Sargazi et al. 2021). For instance, the synthetic compound, chloroquine (CHL), the drug of choice for malaria, has recently gained importance for its off-target activity on GB, wherein Sotelo *et al*., and Briceno *et al*., had reported that addition of CHL as adjuvant to TMZ had significantly prolonged the survival of GB patients and several clinical trials are underway (Briceño et al. 2003; Sotelo et al. 2006). Also, repurposing natural derivatives such as flavonoids and marine compounds had gained increasing interest in GB research (Tseng et al. 2004; Do et al. 2010; Sabarinathan et al. 2011; Lu et al. 2012; Du et al. 2013; Daisy et al. 2021). For instance, naringenin (NAR), a bioactive flavonoid, was reported to alter the mitogen-activated protein kinase (MAPK) signaling pathway in glioma and induce apoptosis via modulation of Bax/BCL-2 ratio (Sabarinathan et al. 2010; Aroui et al. 2016). Further, phloroglucinol (PGL), a marine derivative had induced apoptosis by modulating the endoplasmic reticulum (ER) stress pathway in U-87, U251 and C6 glioma cells (Lu et al. 2012). However, the therapeutic effect of these drugs on sensitizing GB, as monotherapies, are not up to the par. Hence, we were interested to determine the therapeutic effect of the combined administration of these drugs (CHL, NAR, PGL) as a cocktail along with the standard drug (TMZ).

Interestingly, our previous research found that co-administration of these drugs as a quadruple-drug combination (TCNP) had profound effect on hampering malignant glioma proliferation and inducing apoptosis *in vitro* in C6, U-87 MG and LN229 glioma cells and *in vivo* (Daisy et al. 2021; Precilla et al. 2022). Initially, the dual-drug and triple-drug combinations were assessed for their cytotoxicity effects in all the three glioma cell lines (Precilla et al. 2022). Interestingly, we found that individually, all the drugs depicted a dose-dependent reduction in the cellular viability; though the dual-drug and triple-drug treatments depicted a relatively moderate patterns of reduction in cell viability in all the cell lines, yet they were not statistically significant. Nevertheless, administration of these drugs as a quadruple-drug combination (TCNP) had induced cytotoxicity in a synergistic and statistically significant manner than the dual-and triple-drug treatments. As a step forward, these drugs were taken to a preclinical setup, wherein, we observed that the quadruple-combinatorial treatment (TCNP) had induced significant pathological effect and prolonged the overall survival of orthotopic xenografts amidst being less toxic to other vital organs (Precilla et al. 2022).

However, besides observation of their pathological effect, studies that shed molecular insights on the mechanism of action of the drugs would lay a profound background for further investigations on this quadruple-drug combination (TCNP). This is because, being a tumor with complex heterogeneity, GB is characterized by aberrant activation of major signal transduction pathways such as PI3K/AKT/mTOR, RAS/RAF/MAPK, Notch, Hedgehog and WNT signalling pathways which contributes to glioma proliferation (Jeuken et al. 2007; Fan and Weiss 2010; Stockhausen et al. 2010; Liu et al. 2011, 2014; Precilla et al. 2021; Daisy Precilla et al. 2022). Among these, the WNT/ β-catenin signalling pathway is of paramount importance in gliomagenesis, as it enables GB cells to recapitulate the embryonic processes, thereby promoting glioma growth, progression and therapeutic resistance (Latour et al. 2021).

Compelling evidences on the involvement of WNT/β-catenin signalling in malignant glioma have reported that, besides contributing to tumour vasculature, the WNT pathway also contributes to TMZ therapeutic resistance (Yun et al. 2020; Daisy Precilla et al. 2022). Additionally, WNT regulates the expression of epidermal growth factor receptor (EGFR), a key angiogenic factor in various cancers including glioma and is also partially responsible for epithelial mesenchymal transition (EMT) in various cancers (Chen et al. 2012; Paul et al. 2013). Thus, the WNT signalling cascade is an intricate and ubiquitous pathway which influences GB pathogenesis; Therefore, inhibition of WNT/β-catenin signalling pathway may serve as a rational and multipronged target for malignant glioma to suppress tumor proliferation, invasion and surrounding vasculature as an effort to improve the therapeutic outcome of GB patients (McCord et al. 2017). Hence, the current study has been aimed to investigate the implications of these drugs both individually and as a quadruple-drug regime (TCNP) on modulating WNT signalling pathway as a therapeutic target for GB.

## 2. MATERIALS AND METHODS

### 2.1 Reagents

Minimal Essential Medium Eagle (EMEM) (Catalogue No: AT047) and penicillin/streptomycin antibiotic solution were procured from Hi-Media, Mumbai. Fetal Bovine Serum (FBS), Pierce® RIPA buffer, Bicinchoninic acid (BCA) assay kit for protein quantification and MitoProbe JC-1 Assay kit for flow cytometry were procured from Thermo Fisher Scientific, South America. DMSO and the drugs employed for the study namely, TMZ, CHL, NAR and PGL were obtained from Sigma Aldrich, USA. Desalted oligonucleotide primers for EGFR, β-catenin, c-myc, c-jun, ERK-2, BAD, Cytochrome-c (cyt-c), caspase-8, caspase-3, BCL-2 and β-actin were purchased from Bioserve, Telangana, India. PrimeScript RT reagent kit, *in-situ* apoptosis detection kit and TB Green Premix Ex Taq II were procured from Takara Bio Inc, Japan. Polyclonal antibodies against Ki-67 (Cat. No. E-AB-22027), EGFR (Cat. No. E-AB-31281), β-catenin (Cat. No. E-AB-22111), c-myc (Cat. No. E-AB-30975), c-jun (Cat. No. E-AB-30513), ERK-2 (Cat. No. E-AB-31374), BAD (Cat. No. E-AB-60360), β-actin (Cat. No. E-AB-20058) and goat anti-rabbit IgG-HRP secondary antibody (Cat. No. E-AB-AS014) were purchased from Elabscience Biotechnology Inc, Hubei, China. cyt-c (Cat. No: E-EL-H0056), Caspase-8 (Cat. No: E-EL-H0659), Caspase-3 (Cat. No: E-EL-R0160) and BCL-2 (Cat. No: E-EL-H0114) ELISA kits were purchased from Elabscience Biotechnology Inc, Hubei, China. WesternBright ECL substrate kit (Cat. No. K-12045-D20) was obtained from Advansta, San Jose, CA, USA. All the other unspecified reagents were procured from Thermo Fisher Scientific, South America.

### 2.2 Cell Line and culture conditions

U-87 MG human GB cell line (RRID: CVCL_GP63) (Passage Number: 58) was procured from National Centre for Cell Sciences (NCCS) (Pune, India) and authenticated by short tandem repeat (STR) profiling. The tested cell line sample was 100% similar to ATCC STR profile database. The cell line was free from mycoplasma contamination, that was detected by Hoechst staining and PCR. The cells were maintained in EMEM supplemented with 10% FBS and 1% antibiotic solution. The cells were sustained at 37 ºC under humidified atmosphere of 95% air and 5% CO_2_. At their logarithmic growth phase, the cells were trypsinized, counted using trypan blue dye-exclusion method and resuspended in 1X-PBS to a final concentration of 1×10^6^ cells per 3 μl of PBS.

### 2.3 Animals

Healthy male Wistar rats (180-240 grams; 8-10 weeks old, n= 49) were purchased from Biogen Pvt. Ltd. (Bangalore, India) and were maintained under optimum humidity (50-60%), temperature (22±1ºC) and light conditions (12 hours light-12 hours dark cycle), with pellet food and water provided *ad libitum*. All the animal experiments carried out in the study were in compliance with ethical standards and approved by the Institutional Animal Ethics Committee (03/IAEC/MG/09/2021-II). Experiments were conducted based on the Control and Supervision of Experiments on Animals (CPCSEA) and Animal Research: Reporting of In Vivo Experiments (ARRIVE) guidelines and were in accordance with National Research Council’s Guide for the Care and Use of Laboratory Animals.

#### 2.3.1 Intracranial tumor engraftment

To orthotopically implant GB cells into healthy rats, stereotactic surgery was carried out according to the protocol of Baker et al., (Baker et al. 2015). For this, the xenograft recipient was sedated using ketamine and xylazine cocktail (87 mg/kg and 13 mg/kg body weight). The skull was exposed by incising the skin from the frontal to the occipital bone. After marking the implantation location (3.00 mm lateral and 0.5 mm posterior position), a small burr hole was drilled using a cordless power drill. Followingly, a total of 1×10^6^ U-87 MG GB cells suspended in 3 μl PBS was delivered into the target site at a depth of 6 mm. The burr hole was then sealed with bone wax and appropriate post-operative relief measures were carried out.

#### 2.3.2 Animal Grouping

20 days post-implantation, the animals were randomly divided into six groups with 7 animals in each group as follows (**Fig. S1**). The drug doses for various experimental groups employed for the current study was determined based on the previous literature (Golden et al. 2014; Fernandes et al. 2017; Pranav Nayak et al. 2019; Seung et al. 2021; Shi et al. 2021).

**Group 1 (Control group, n=7):** Healthy animals without tumor implantation or drug treatment.

**Group 2 (Tumor-Control group, n=7):** Untreated animals injected with U-87 MG GB cell suspension (1×10^6^ cells in 3 μl PBS) alone.

**Group 3 (TMZ group, n=7):** Animals administered with 10 mg/kg body weight of TMZ dissolved in 0.1% DMSO every day.

**Group 4 (CHL group, n=7):** Animals administered with 10 mg/kg body weight of CHL dissolved in saline every day.

**Group 5 (NAR group, n=7):** Animals administered with 10 mg/kg body weight of NAR dissolved in 0.1% DMSO every day.

**Group 6 (PGL group, n=7):** Animals administered with 50 mg/kg body weight of PGL dissolved in 0.1% DMSO every day.

**Group 7 (TCNP group, n=7):** Animals administered with the combination of all the four drugs, TMZ (10 mg/kg/day) + CHL (10 mg/kg/day) +NAR (10 mg/kg/day) +PGL (50 mg/kg/day) dissolved in vehicle every day.

All the drugs were administered orally for 14 days. At the end of the treatment period, the rats from each experimental group were euthanized, brains were decapitated and stored in formalin or liquid nitrogen for further analysis. Additionally, blood samples from each experimental group were collected by cardiac puncture method. The supernatant containing plasma samples was collected after centrifugation at 2,500 rpm at room temperature for 10 minutes and was stored at -80 ºC for ESI-Q-TOF-MS analysis.

### 2.4 Immunohistochemistry Analysis

To determine the pathological effects of the drugs on GB xenografts, immunohistochemistry (IHC) analysis was performed. For this purpose, tissue sections from each experimental group were formalin-fixed and sectioned (5 μm). Following deparaffinization, antigen retrieval was performed by heating the formalin-fixed sections in microwave for 20 minutes in sodium citrate buffer containing 10 mM Sodium citrate and 0.05% Tween-20 (pH 6.0). Endogenous peroxidase activity was blocked by incubating the tissues in 3% H_2_O_2_ for 15 minutes. The tissue sections were then blocked with 3% bovine serum albumin (BSA) for 10 minutes. After washing in 1X-TBST buffer twice, the slides were sequentially incubated with primary antibodies (rabbit anti-Ki67 1:200 dilution; rabbit anti-EGFR antibody 1:200 dilution; rabbit anti-β-catenin antibody 1:200 dilution; rabbit anti-BAD antibody 1:200 dilution) for 1 hour at room temperature; After washing off the unbound antibodies, the sections were then incubated with HRP-conjugated secondary antibody for 30 minutes at room temperature. The bound antibodies were detected using diaminobenzidine (DAB), counterstained with haematoxylin and mounted using mounting medium (Dako, Denmark). Microscopic images were obtained using inverted microscope (Primo Star, Carl Zeiss, Germany). Quantification of the acquired images was performed using Fiji software (Yang et al. 2021).

### 2.5 Quantitative Real-Time PCR Analysis

To assess the relative gene expression levels of WNT signalling and apoptotic markers in the GB-induced and drug-treated xenografts, qRT-PCR analysis was carried out according to the standard protocol. Briefly, total RNA was extracted from the tissue samples by TRIzol reagent (Invitrogen, CA, USA) and reverse transcribed using PrimeScript RT Reagent kit (Takara Bio Inc, Japan). For qRT-PCR, the complementary DNA (cDNA) products were amplified using TB Green Premix Ex Taq II (Takara Bio Inc, Japan) according to the manufacturer’s instructions in CFX 96 thermo cycler (Bio-Rad Laboratories, Hercules, CA, USA). The thermal cycling conditions were as follows: initial denaturation at 95ºC for 30 seconds followed by 39 cycles of 95ºC for 5 seconds, 60ºC for 30 seconds, 72ºC for 30 seconds and a final extension of 72ºC for 3 minutes. The 2^-ΔΔCq^ method was used to calculate the fold change value. β-actin was used as a normalizer gene. All the primers used for the current study are listed in **Table S1**.

### 2.6 Immunoblotting Analysis

To assess the protein expression levels of WNT signalling and apoptotic markers in the GB-induced and drug-treated xenografts, western blotting was carried out. Briefly, protein lysates were extracted from the brain tissues of various experimental groups using Pierce® RIPA lysis and extraction buffer with protease and phosphatase inhibitor cocktail. Following incubation for 30 minutes on ice, the lysates were centrifuged at 12,000 g for 30 minutes at 4ºC. The supernatant containing protein lysates were quantified using BCA assay kit and equal amounts of protein (30 μg) were subjected to SDS-PAGE. Separated proteins were then transferred to polyvinylidene difluoride (PVDF) membranes (0.45 μM) and blocked with 3% BSA for 30 minutes. The membranes were then probed overnight with the primary antibodies (1:1000 dilution); β-actin was used as an internal reference standard. Following wash with IX-TBST buffer, the membranes were then incubated with secondary antibody (goat anti-rabbit IgG-HRP, 1:10,000 dilution) at room temperature and developed using WesternBright ECL kit. Chemiluminescence was detected employing ChemiDoc™ MP Imaging System (Bio-Rad Laboratories Inc, Hercules, CA, USA) and the band intensities were quantified using Image Lab software (Bio-Rad Laboratories Inc, Hercules, CA, USA).

### 2.7 ELISA

To detect the protein expression levels of apoptotic markers namely cyt-c, caspase-8, caspase-3 and BCL-2, ELISA was performed according to the manufacturer’s protocol. Briefly, the protein lysates obtained from various experimental groups were loaded in the 96-well plates pre-coated with the monoclonal antibodies (cyt-c, caspase-8, caspase-3 and BCL-2) and incubated for 90 minutes at 37ºC. After removal of the unbound samples by wash buffer, biotinylated antibody working solution was added to each well and incubated for 60 minutes at 37ºC. Later, HRP-conjugated streptavidin was added to all the wells and the solution was incubated for 60 minutes at 37ºC. Followingly, 90 μl of substrate was added to each well (incubation at 37ºC for 30 minutes). To stop the reaction, 50 μl of stop solution was added to each well and the absorbance was measured at 450 nm immediately using spectrophotometer (Molecular Devices Spectra-Max M5, USA). Relative protein concentrations were then determined by interpolating the absorbance with that of the standard curve obtained from the standard protein.

### 2.8 Flow cytometric detection of mitochondrial membrane depolarization (ΔΨm)

To determine the changes in the ΔΨm by flow cytometry, brain tissues from each experimental group were stained using MitoProbe JC-1 assay kit according to the manufacturer’s protocol. Briefly, the astrocytes were isolated from the brain tissues of each experimental group in accords with the protocol of Schildge et al., (Schildge et al. 2013). Isolated astrocytes were washed in 1X-PBS twice and incubated with JC-1 staining solution for 30 minutes in dark. Cell pellet obtained after centrifugation was then subjected to flow cytometry (BD FACSCelesta™, BD Biosciences, San Jose, CA, USA). Totally 30,000 gated events were analysed for each group with emissions set at 590 nm and 530 nm for red and green fluorescence respectively. Data analysis was done using FlowJo Software (FlowJo ™, BD Biosciences, USA).

### 2.9 TUNEL Assay

To detect the induction of apoptosis in drug-treated GB xenografts, TUNEL assay was carried out using *in-situ* apoptosis detection kit (Takara Bio Inc, Japan), as per the manufacturer’s protocol. Briefly, 5 μm paraffin embedded tissue sections were de-waxed, and incubated with Proteinase-K (20 μg/ml) at room temperature for 15 minutes. Endogenous peroxidase activity was blocked by incubating the sections with 3% H_2_O_2_ for 15 minutes. Followingly, the slides were exposed to the TUNEL labelling reaction mixture consisting of DIG-dUTP, reaction buffer and terminal deoxynucleotidyl transferase for 90 minutes at 37ºC in a humidified chamber. The reaction was then terminated by washing the slides in 1X-PBS. Subsequently, the tissue sections were incubated with 70 μl of anti-FITC HRP conjugate for 30 minutes followed by colouring with DAB and counterstaining with haematoxylin. The brown coloured TUNEL-positive cells were then visualized by inverted microscopy (Primo Star, Carl Zeiss, Germany). The number of apoptotic cells were determined by dividing the number of apoptotic cells by the total number of cells in the field (Garrity et al. 2003).

### 2.10 ESI-Q-TOF MS Analysis of the drugs and their metabolites

In order to elucidate if the chosen drugs both individually and as a combination (quadruple-drug) had breached the blood-brain barrier (BBB) and reached the target site to exert their pharmacological effects, ESI-Q-TOF MS analysis was performed employing the brain tissues and plasma samples obtained from various experimental treatment groups. For this purpose, the samples were processed as follows:

### 2.10.1 Sample Preparation

Approximately 500 mg of frozen brain tissues were homogenized in 500 μl of distilled water. After centrifugation at 2,500 rpm for 10 minutes, the supernatant containing tissue lysates was collected and transferred to a fresh tube. 100 μl of these tissue lysate aliquots and rat plasma samples, separately, were centrifuged at 3,000 rpm for 10 minutes. Following centrifugation, 5 μl of the tissue and plasma supernatants from each experimental group was diluted with methanol and injected directly into the ion source.

### 2.10.2 ESI-Q-TOF MS conditions

Drug detection studies were performed using Accurate Mass-Q-TOF mass spectrometer (Agilent Technologies Inc, USA) equipped with electron spray ion source (ESI), operated in positive mode. The mobile phase consisted of acetonitrile and water containing 0.01% formic acid (1:1). Analyses were carried out at a flow rate of 0.1 ml per minute.

### 2.11 Statistical Analysis

All the experiments were conducted in triplicates. Analysis of the experimental data was carried out using GraphPad Prism v.8.0.1 (GraphPad Software, San Diego, CA, USA). Results were represented as mean ± standard deviation (SD). Statistical analysis between the control and various experimental groups was performed using Analysis of Variance (ANOVA). Post-hoc analysis was carried out using Dunnett or Tukey’s post-hoc test. p < 0.05 was considered to be significant.

## 3. RESULTS

### 3.1 Quadruple-combinatorial treatment had reduced tumor proliferation at morphological level in GB xenografts

Given the evidence that the quadruple-combinatorial treatment (TCNP) had suppressed GB growth and proliferation *in vitro* and prolonged the overall survival rate with significant pathological effect on tumor tissues *in vivo* in our previous study (Precilla et al. 2022), we were interested to investigate the underlying *in vivo* efficacy of this combination in a broader aspect. To accomplish this, initially, IHC staining was performed on the tissue sections of GB-induced and drug-treated xenografts (individual and combinatorial-drug) against the markers of GB proliferation (Ki-67, EGFR), WNT signaling (β-catenin) and apoptosis (BAD). Interestingly, we observed that the quadruple-combinatorial treatment (TCNP) had significantly reduced the expression of the proliferative markers namely, Ki-67 (3.5%) (**Fig. 1- A- green arrowheads**) and EGFR (11%) (p<0.001) (**Fig. 1- B- green arrowheads**) in comparison with the tumor-control (**Fig. 1- A, B- red arrowheads**) and individual drug-treated sections (**Fig. 1-A, B- red arrowheads**). Furthermore, the combinatorial treatment (TCNP) was also bestowed with the potency to suppress the expression β-catenin (5%) more significantly (p<0.001) (**Fig. 1- C- green arrowheads**), than the individual regimes (**Fig. 1- C- red arrowheads**). Parallelly, while suppressing GB proliferation and WNT signaling, the quadruple-combinatorial treatment (TCNP) had significantly facilitated apoptosis as evident from positive-staining results for BAD (87%) (p<0.001), a pro-apoptotic marker (**Fig. 1- D- red arrowheads**) when compared with the tumor control (**Fig. 1- D- green arrowheads**). While individually, TMZ, CHL, NAR and PGL had depicted modest effect on suppressing GB proliferation, WNT signaling and inducing apoptosis, they were not as efficient as the quadruple-drug combination (TCNP) (**Fig. 1- A- D- red arrowheads**). Taken together, our IHC results indicated that the quadruple-combinatorial treatment (TCNP) could suppress GB tumor proliferation at morphological level.

**Fig. 1.**
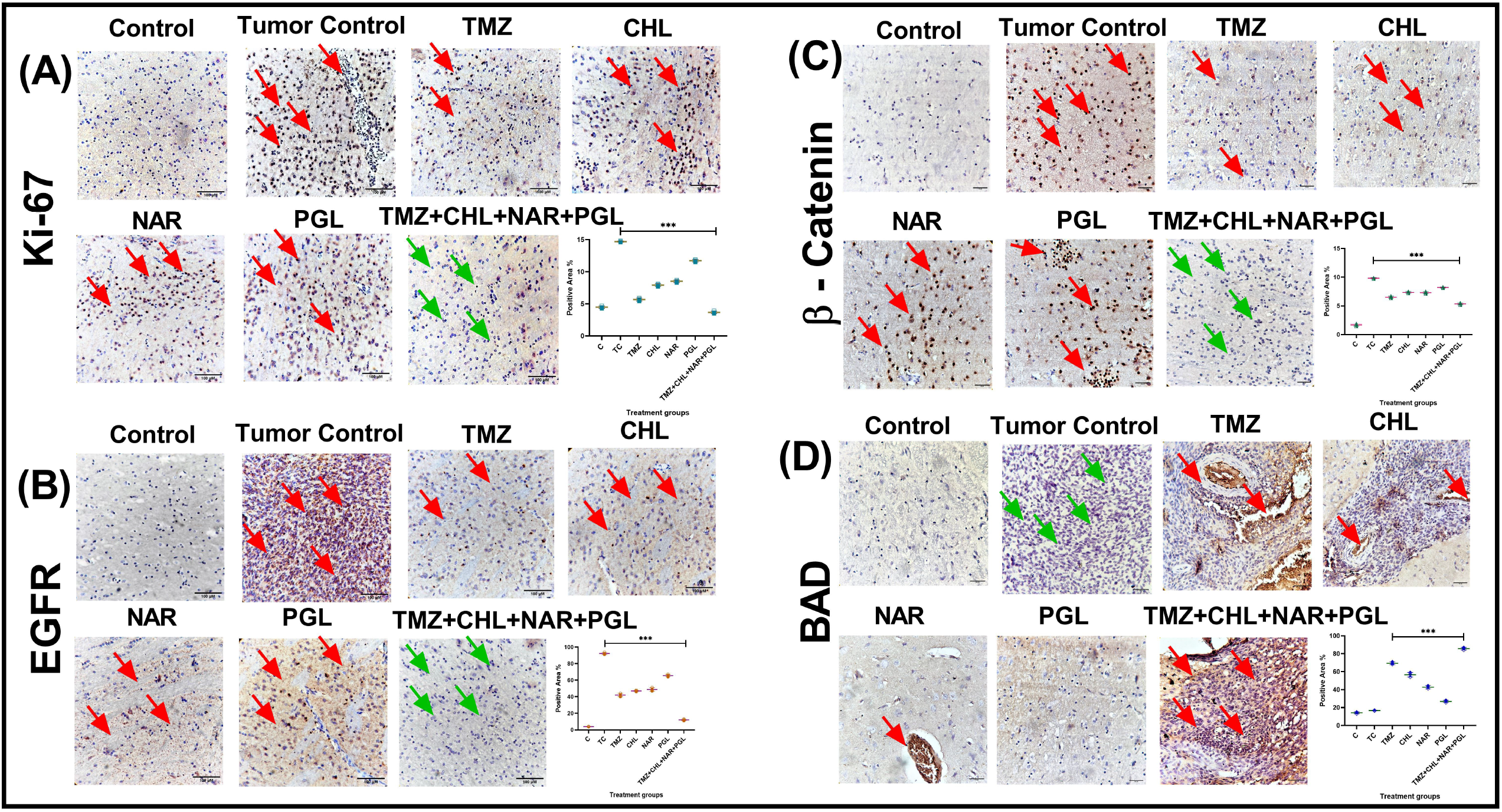
Effect of the quadruple-combinatorial treatment on the key markers of GB proliferation and apoptosis in GB-bearing xenografts. **(A-D)** Patterns of positive/negativeimmunostaining for Ki-67, EGFR, β-catenin and BAD following administration of the individual and combinatorial treatments were identified at 40x in inverted light microscope (positive staining-red arrowheads, negative staining-brown arrowheads). Quantitative data of the staining results were obtained using Fiji software, ***p<0.001 in comparison with the tumor control, n=3

### 3.2 Quadruple-combinatorial treatment induced apoptosis by modulating WNT/ β-catenin signaling pathway in GB xenografts at transcriptional level

Following assessment of the anti-proliferative efficacy the quadruple-combinatorial treatment (TCNP), we were interested to unravel their underlying molecular mechanism on GB xenografts. From our IHC findings, we observed a decreased expression of β-catenin in the quadruple-drug treated rats (**Fig. 1- C- green arrowheads**), with prominent anti-proliferative efficacy against Ki-67 and EGFR (**Fig. 1- A, B- green arrowheads**). Based on this result, we hypothesized that this effect would have been mediated by the inhibition of WNT/β-catenin signaling pathway. Nevertheless, previous studies have shown that WNT/β-catenin signaling is aberrantly activated in glioma and contributes to GB tumorigenesis (Zhang et al. 2012). Importantly, studies have suggested that regulation of WNT/β-catenin signaling using chemotherapeutics had depicted promising effects on inhibiting tumor growth, proliferation and inducing apoptosis (Park et al. 2013; Li et al. 2017; Wu et al. 2018).

Thereof, in order to explore the regulatory mechanism of our quadruple-drug combination (TCNP) on WNT/β-catenin signaling and apoptosis induction, the mRNA expression profiles of WNT signaling and apoptotic markers were assessed by qRT-PCR analysis. Interestingly, our results showed that the combinatorial treatment (TCNP) could efficiently modulate WNT signaling pathway *in vivo* by reducing the levels of EGFR (**Fig. 2- I- A-C**) (p<0.001) and the key components of WNT signaling. Particularly, when compared with the tumor control, β-catenin and its downstream oncogenes namely, c-myc, c-jun and ERK-2 were significantly down-regulated (p<0.001) following administration of the quadruple-combinatorial treatment (TCNP) **(Fig. 2- I- A, C**). Individually, treatment with TMZ had down-regulated WNT/β-catenin pathway genes moderately but was not statistically significant (**Fig. 2- I- A, C**). However, CHL, NAR and PGL, as individual agents showed relatively modest activity on modulating WNT/β-catenin signaling pathway *in vivo* (**Fig. 2- I- A, C**).

**Fig. 2.**
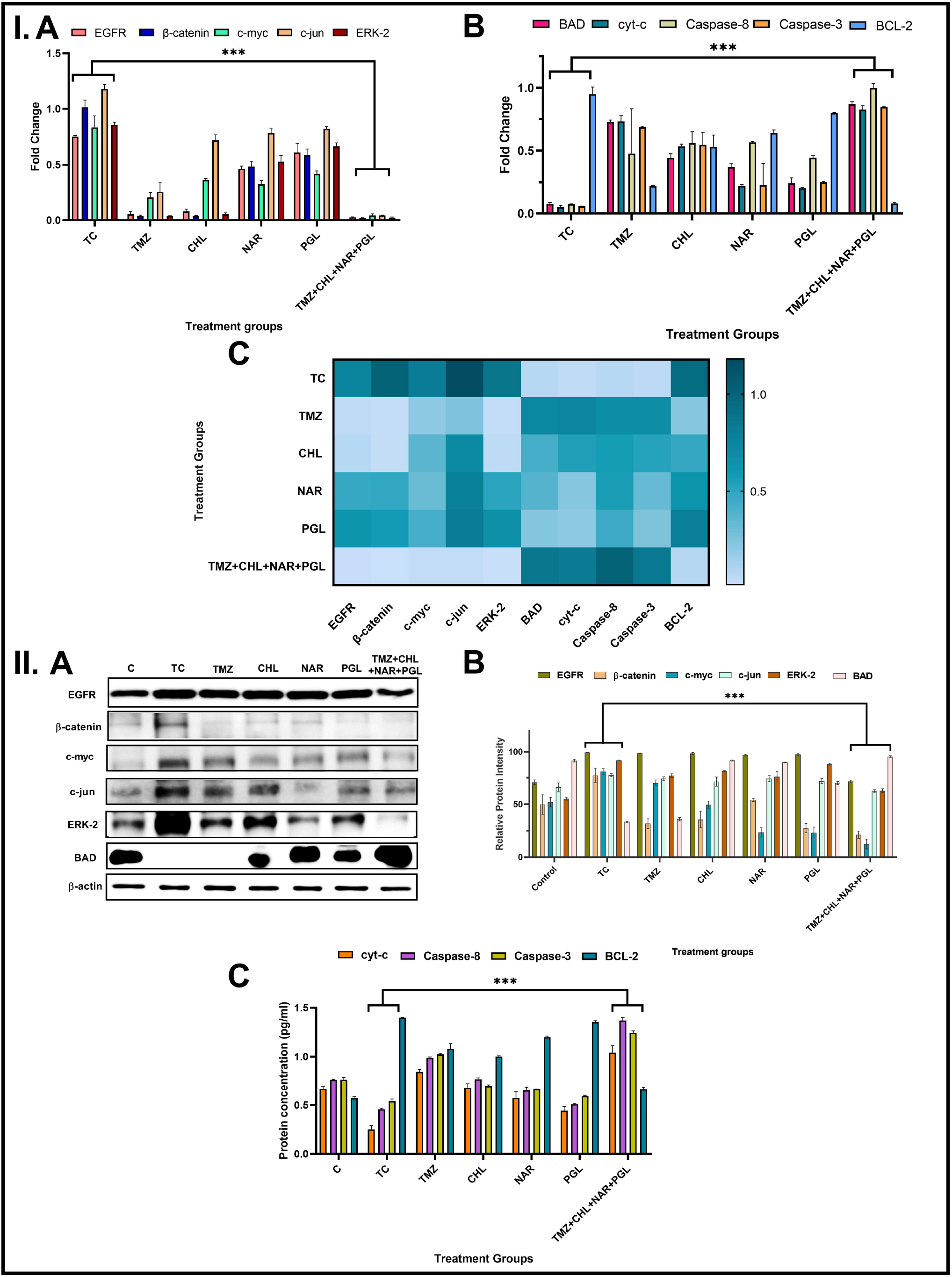
Effect of the quadruple-combinatorial treatment on modulation of WNT/β-catenin signaling and induction of apoptosis in GB-bearing xenografts. **(I.)** Brain tissues of the GB-bearing xenografts, following administration of the individual and combinatorial treatments were analysed for the gene expression levels of WNT/ β-catenin **(A, C)** and apoptotic markers by qRT-PCR **(B, C). (II)** Brain tissues of the GB-bearing xenografts, following administration of the individual and combinatorial treatments were analysed for the protein expression levels of WNT/ β-catenin and apoptotic markers by Western Blotting **(A, B)** and ELISA **(C)**. Quantitative data of the immunoblot bands were obtained using Image Lab software. Data are represented as Mean ± SD, ***p<0.001 in comparison with the tumor control, n=3

Accumulating evidences had suggested that down-regulation of WNT/β-catenin signaling results in apoptosis in various cancers (Jiang et al. 2013; Lan et al. 2015). Hence, subsequently, we validated the expression levels of apoptotic markers by qRT-PCR. Interestingly, we observed that treatment with the quadruple-drug combination (TCNP) had significantly elevated the levels of cyt-c, BAD, caspase-8, and caspase-3 (p<0.001) while the levels of BCL-2, an anti-apoptotic gene was significantly down-regulated (p < 0.001) (**Fig. 2- I- B, C**). Individually, all the chosen drugs did depict a relatively moderate apoptotic effect, but were not as significant as the quadruple-combinatorial treatment (**Fig. 2- I- B, C**). Overall, our gene expression results indicated that the combinatorial treatment (TCNP) could significantly down-regulate WNT/β-catenin signaling cascade and induce apoptosis more effectively than the individual counterparts.

### 3.3 Quadruple-combinatorial treatment induced apoptosis by modulating WNT/ β-catenin signaling pathway in GB xenografts at translational level

To elucidate if the patterns of gene expression profiles of WNT/β-catenin signaling and apoptotic markers was also reconciled at the protein expression level, immunoblotting and ELISA was performed. Interestingly, the quadruple-combinatorial treatment (TCNP) had significantly reduced the expression of EGFR, β-catenin and its downstream targets namely, c-myc, c-jun, and ERK-2 than their individual counterparts (p<0.001) (**Fig. 2- II- A, B**). Followingly, the protein expression levels of apoptotic markers were assessed whereby it was evident that the quadruple-drug treatment (TCNP) had significantly induced apoptosis in the orthotopic GB xenografts by activating the intrinsic apoptotic cascade via upregulation of cyt-c, BAD, caspase-8 and caspase-3 with concomitant downregulation of BCL-2 (p<0.001) (**Fig. 2- II- A- C**). Individually, all the chosen drugs had depicted moderate activity on downregulating the key drivers of the WNT/β-catenin signaling cascade and promoted apoptosis but were not statistically significant when compared with the quadruple-combinatorial treatment (TCNP) (**Fig. 2- II- A- C**). Taken together, our mechanistic study emphasized that the anti-GB efficacy of the quadruple-combinatorial treatment (TCNP) was mediated via the modulation of the WNT/β-catenin signaling cascade, thereby inducing apoptosis in the orthotopic GB xenografts.

### 3.4 Quadruple-combinatorial treatment triggered mitochondrial depolarization (ΔΨM) in GB xenografts

Following affirmation of the anti-proliferative and apoptotic effects of the quadruple-drug combination (TCNP) by various staining and mechanistic studies, we wanted to investigate if the apoptotic effect was mediated by the involvement of mitochondria, as the levels of cyt-c and BAD were significantly upregulated in the quadruple-drug treated GB xenografts. For this purpose, the alterations in mitochondrial membrane potential (MMP), was assessed (Jo et al. 2005). The red/green fluorescence ratio of JC-1 dye can be used to directly assess the alterations in MMP. Healthy cells exhibit a higher red fluorescence, indicating higher MMP activity while apoptotic cells exhibit more green fluorescence and low MMP (Sivandzade et al. 2019). Interestingly, we found that the quadruple-combinatorial treatment (TCNP) had significantly altered the MMP, as evident from the increase in the green fluorescence (80%) (p<0.001), when compared with the tumor control and other treatment groups (**Fig. 3- A**). Among the individual drugs, PGL and TMZ depicted moderate alterations in MMP (17% & 10.1% of green fluorescence) but they were not statistically significant (**Fig. 3- A**). However, as individual regimes, CHL and NAR did not depict any effects on mitochondrial depolarization (**Fig. 3- A**). Since the loss of MMP is an early indicator of apoptosis, our results suggest that the quadruple-combinatorial treatment (TCNP) could effectively promote apoptosis by altering the MMP in GB xenografts.

**Fig. 3.**
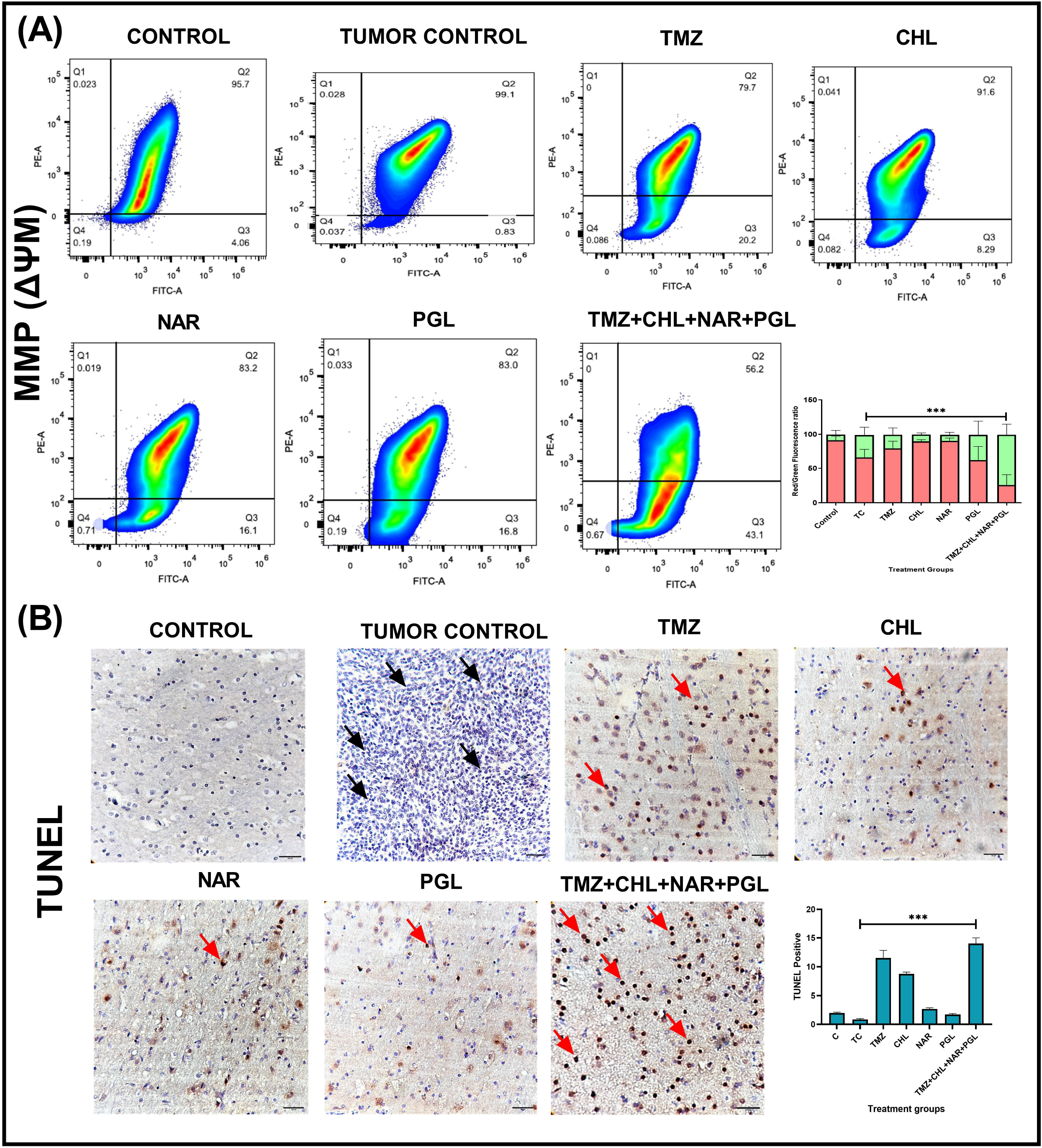
Effect of the quadruple-combinatorial treatment on induction of apoptosis and alteration of MMP (ΔΨM) in GB-bearing xenografts. **(A)** Patterns of shift in the red/green fluorescence following administration of the individual and combinatorial treatments were quantified by flow cytometry. **(B)** Patterns of positive/negative TUNEL staining following administration of the individual and combinatorial treatments were identified at 40x in inverted light microscope (positive staining-red arrowheads, negative staining-brown arrowheads). Quantitative data of the staining results were obtained using Fiji and FlowJo software, ***p<0.001 in comparison with the tumor control, n=3

### 3.5 Quadruple-combinatorial treatment induced apoptosis in GB xenografts

To substantiate the efficacy of the quadruple-combinatorial treatment (TCNP) on inducing apoptosis following our immunostaining and expression studies at transcriptional and translational level, the tissue sections from each experimental group were subjected to TUNEL assay. Interestingly, we found that the highest percentage of apoptotic cells (15%) was observed in the quadruple-drug treated (TCNP) sections **(Fig. 3- B- red arrowheads**) in comparison with the tumor control (0.7%) (**Fig. 3- B- black arrowheads**), wherein major proportion of cells exhibited shrunken brown stained nuclei (p<0.001). Individually, TMZ depicted a moderate percentage of apoptotic cells (10%), but was not statistically significant (**Fig. 3- B- red arrowheads**). Though CHL, NAR and PGL displayed few apoptotic cells (8.7%, 2.8% & 1.8%), their activity as individual regimes were not effective when compared to their efficacy as a quadruple-combinatorial treatment (**Fig. 3- B- red arrowheads**). Thus, our TUNEL staining results strongly indicated that the quadruple-drug combination (TCNP), beside restraining GB proliferation via modulating WNT/β-catenin signaling cascade, could effectively induce apoptosis in the *in vivo* experimental GB model.

### 3.6 Quadruple-combinatorial treatment was detected in the brain tissue and plasma samples of GB xenografts

One of the key obstacles for treatment of malignant gliomas is the delivery of chemotherapeutic agents across the BBB (Zhan and Lu 2012). Hence, to determine if the systemically delivered quadruple-drug (TCNP) had the ability to cross the BBB and elicit its therapeutic efficacy, we investigated if the drugs and their metabolites, both individually and in combination were detected in the brain tissues and plasma of the GB-bearing drug-treated xenografts. To test this, the brain tissue and plasma samples of the drug-treated GB xenografts were subjected to ESI-Q-TOF MS analysis in positive ion mode. Interestingly, we found that all the chosen drugs had breached the BBB and were detected in the brain tissue and plasma samples as well (**Fig. 4, Fig. S2**). When administered as a quadruple-drug combination (TCNP), in the rat plasma and brain tissue samples, TMZ (C_6_H_6_N_6_O_2_, m/z= 194.151) was detected at [M^+^Na]^+^-217.04 and [M^+^H]^+^-195.06, while its metabolite 4-amino-5-imidazole-carboxamide (AIC) (m/z= 126.12) was detected at [M^+^H]^+^-127.06 (**Fig. 4**). Parallelly, CHL (C_18_H_26_CIN_3_, m/z= 320.0) was detected in the form of its active metabolite desethylchloroquine (C_16_H_22_CIN_3_, m/z=292.2) at [M^+^H] ^+^ - 292.16 (**Fig. 4**); NAR (C_15_H_12_O_5_, m/z= 272.25) and PGL (C_6_H_6_O_3_, m/z= 125), on the other hand, showed molecular ion peaks at [M^+^H] ^+^ - 273.07 and 127.03 respectively (**Fig. 4**). Nevertheless, as individual regimes, all the drugs were detected in the plasma and brain tissues of the individual drug treated GB xenografts; Among them, TMZ was detected as both parent ion at [M^+^H] ^+^ - 195.06 and as its active metabolite, AIC at [M^+^H] ^+^- 127.06 (**Fig. S2**). Similarly, CHL was detected as a parent ion at [M^+^H] ^+^ - 320.18 and as its metabolite, desethylchloroquine at [M^+^H] ^+^ - 292.15 (**Fig. S2**); Albeit, NAR and PGL were detected at [M^+^H] ^+^ - 273.07 and 127.03 respectively (**Fig. S2**). Thus, our ESI-Q-TOF MS data suggests that, the systemic delivery of the drugs as a quadruple-drug cocktail could successfully breach the BBB and reach the target site to elicit therapeutic response in a qualitative manner.

**Fig. 4.**
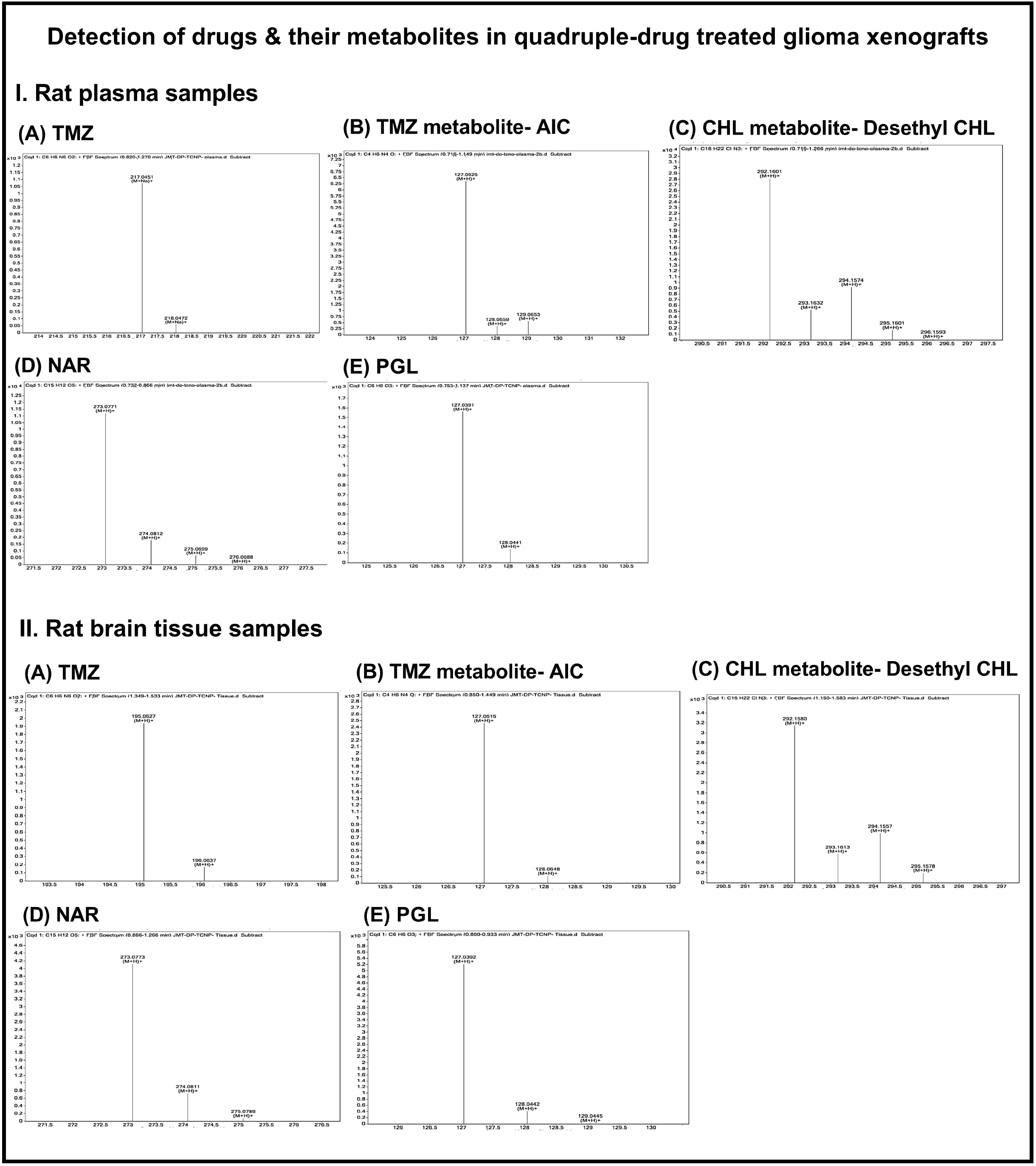
Ability of the quadruple-combinatorial treatment to breach the BBB and exert its therapeutic effect in GB-bearing xenografts. Drug detection was assessed by ESI-Q-TOF MS in the GB-bearing-combinatorial drug-treated rat brain tissues and plasma samples

## 4. DISCUSSION

Malignant gliomas are intrinsic brain tumors accounting for 81% of the CNS malignancies (Iuchi et al. 2022). Despite the cutting-edge treatment modality comprising surgery, radio- and chemotherapy, long-term control over GB progression remains elusive and tumor recurrence is inevitable (van Linde et al. 2017). Thus, there exists an urgent need for effective alternate therapeutic strategies to combat GB pathogenicity.

In this context, drug repurposing would serve as an attractive tactic as the pharmacological profiles of chosen drugs are well-known which will reduce the cost investment and can be tested in very small cohorts as well (Cha et al. 2018, Padhy and Gupta 2011). Nevertheless, the process of identifying and validating the potent candidates for repurposing ranges from experimental (*in vitro* and *in vivo*) studies to randomized clinical trials (Saeidnia et al. 2015). While considering drug repurposing for malignancies like glioma, the therapeutic efficacy of the drugs as monotherapies seems to be less efficient (Ho and Ho 2021); thereof, the focus has now shifted to identify synergistic combination therapies employing two or more drugs (Brown et al. 2008; Kuduvalli et al. 2021; Daisy et al. 2021). In line with this concept, herein, we have presented the *in vivo* evidence for GB treatment enhancement using a quadruple- combinatorial drug treatment (TCNP) comprising a cocktail of both synthetic and natural derivatives.

This study was designed on grounds of our previous findings, whereby it was evident that the quadruple-drug treatment (TCNP) was bestowed with promising anti-glioma efficacy *in vitro* than the dual and triple-drug treatments (Precilla et al. 2022). However, streamlining the molecular mechanism behind this anti-glioma efficacy requires profound investigation. Moreover, it is not surprising that the molecular understanding on GB has undergone a sea change over the last decade. Hence, the current study was designed to gain deep insights into the molecular mechanisms of this quadruple-combinatorial (TCNP) treatment on orthotopic GB xenografts following the preliminary evidences.

In order to provide a more accurate prediction on the clinical outcome of a chemotherapeutic regime, an ideal experimental model is a mandate need. With regard to malignant gliomas, an orthotopic xenograft model is an essential prerequisite, as it would bear the genetic similarities of human glioma and would be highly reproducible (Ji et al. 2016). An interesting takeaway from our study was that such an orthotopic GB model was successfully established (Grade IV glioma). This can be evident from our IHC results, whereby the tumor control group had depicted significant positivity for Ki-67, a key hallmark of GB proliferation (**Fig.1-A- red arrowheads**). Nevertheless, treatment with the drugs as a quadruple-drug combination (TCNP) had significantly reduced the expression levels of Ki-67 (**Fig.1-A- green arrowheads**). Previous studies had shown that individually, TMZ, CHL, and NAR had decreased the levels of Ki-67 in various cancers including glioma (Yang et al. 2011; Zou et al. 2013; Rao et al. 2017; Lin et al. 2018; Choi et al. 2020; Chen et al. 2021). Consistent with those findings, in our study, these drugs had demonstrated a modest activity on reducing the levels of Ki-67 (**Fig. 1- A- red arrowheads**), indicating that their therapeutic efficacy as individual agents were not up to the par. We assume that combined administration of the drugs, CHL, NAR and PGL would have enhanced the efficacy of the benchmark drug, TMZ with a sharp decrease in the levels of Ki-67 (**Fig. 1- A green arrowheads**).

Interestingly, from our preliminary *in silico* analysis, we had observed that all the four drugs namely, TMZ, CHL, NAR and PGL had depicted a profound interaction with two major receptors of WNT/β-catenin signaling pathway namely, Frizzled (FZD) and the co-receptor, the low-density lipoprotein receptor-related protein/alpha 2-macroglobulin receptors (LRP5/6) (Precilla et al. 2021). On grounds of this *in silico* finding, we analysed the regulation of WNT/β-catenin pathway by this quadruple-drug cocktail (TCNP) with major focus on the expression signatures of the main components namely, EGFR, β-catenin, c-myc, c-jun and ERK- 2. Upon activation, by binding of the WNT ligands to FZD receptor and LRP5/6 co-receptor, cytoplasmic β-catenin can translocate into the nucleus, whereby, it binds to TCF/LEF to stimulate the transcription of downstream target genes and inhibit apoptosis (MacDonald et al. 2009). In this context, our immunostaining, transcriptional and translational studies had demonstrated that the administration of the quadruple-combinatorial treatment (TCNP) had suppressed the expression of EGFR and β-catenin (**Fig. 1- B, C- green arrowheads, Fig. 2- I & II**). As the nuclear levels of β-catenin were depleted, this could have prevented the activation of TCF/LEF, which had resulted in the reduction of β-catenin’s downstream targets namely, c-myc, c-jun and ERK-2 (**Fig. 2- I & II**).

Further, as a stepping stone to streamline the down-stream events being modulated following the suppression of WNT/β-catenin signaling, we then studied the expression profiles of the key drivers of apoptosis. Previous studies have reported that when WNT/β-catenin cascade was blocked, the possibility of apoptotic cell death was increased (Wu et al. 2014). As such, under homeostatic conditions, apoptosis can be activated either intrinsically (mitochondria-dependent) or extrinsically (death-receptor mediated) (Kashyap et al. 2021). Both these pathways involve caspases, a family of cystine proteases for activation, that includes the initiator caspases (caspase- 2, 8, 9, 10, 11, 12) and the executioner caspases (caspase- 3, 6, 7) (Prokhorova et al. 2018). In this regard, it is obvious that cell stress or DNA damage often activates apoptosis intrinsically (Wang et al. 2017). As our IHC assessment portrayed that this quadruple-combination (TCNP) had elevated the levels of BAD (**Fig. 1- D- red arrowheads**), a pro-apoptotic protein, we were interested to specifically focus on the intrinsic pathway. Among the key drivers of intrinsic apoptotic pathway, caspase-3 plays a crucial role in inducing apoptosis in a mitochondria-dependent (BCL-2/BAD) manner (Singh et al. 2019). Interestingly, our quadruple-combinatorial treatment (TCNP) had significantly downregulated the levels of BCL-2 and upregulated the expression levels of cyt-c and BAD (**Fig. 2- I & II**). Since BCL-2 is downregulated in the mitochondria, this would have resulted in increased mitochondrial permeability and release of cyt-c. We assume that the released cyt-c would have activated the caspases-cascade, as the expression levels of caspase-8 and caspase-3 were significantly upregulated in the quadruple-drug treated groups (**Fig. 2- I & II**). Thus, our molecular studies emphasized that the quadruple-combinatorial treatment (TCNP) had modulated WNT/β-catenin signaling cascade by induction of apoptosis intrinsically.

As the quadruple-combinatorial treatment (TCNP) elevated the expression levels of cyt c, we were specifically interested to determine if the induction of apoptosis had occurred with the involvement of the mitochondria. This is due to the fact that major events of apoptosis occur in the mitochondria, among which the most significant is the loss of MMP (ΔΨM) (Christmann 2015). Onset of apoptosis is usually associated with the opening of the pores in the mitochondrial membrane which results in the loss of the electrochemical gradient (Burke 2017). Interestingly, we observed that the quadruple cocktail (TCNP) had significantly induced mitochondrial depolarization as evident from our flow cytometric analysis (**Fig. 3- A**). This could be due to the treatment with the quadruple-drug combination (TCNP) that had rendered apoptotic cells with less electronegative mitochondria. This was evident from the high intensity of green fluorescence, as the lipophilic and fluorescent cationic dye, JC-1 was not able to convert into JC-aggregates (red fluorescence) and had remained in the monomeric state (Sivandzade et al. 2019). Contrarily, exposure to the drugs TMZ, CHL, NAR and PGL as individual regimes had rendered a relatively low number of apoptotic cells with the accumulation of JC-1 dye as aggregates with high percentage of red fluorescence than the quadruple-combinatorial treatment (TCNP) (**Fig. 3- A**). This finding was consistent with the results obtained from our *in vitro* study (Precilla et al. 2022). One possible reason for the alteration in MMP following administration of this quadruple-drug combination (TCNP) could be the suppression of the anti-apoptotic protein BCL-2, as it is often expressed in the mitochondria of healthy cells. Thus, from our IHC, molecular and MMP findings, we conclude that the apoptosis induced by this quadruple-combinatorial treatment (TCNP) had involved mitochondria as well.

It is well-known that in tumor cells undergoing apoptosis, the DNA is cleaved at the site of attachment resulting in DNA fragments with free 3’-OH end (Elmore 2007). This facilitates identification of apoptotic cells by transfer of fluorescent-labelled dUTP to the free 3’-OH ends using the enzyme terminal deoxynucleotidyl transferase (TdT) (Ferreira and Afreen 2017). Interestingly, our TUNEL results had depicted that the quadruple-combinatorial treatment (TCNP) had significantly induced apoptosis as evident from the highest percentage of TUNEL-positive cells with brown and shrunken nuclei (**Fig. 3- B- red arrowheads**). While our molecular studies denoted that the quadruple-drug treatment (TCNP) had significantly induced apoptosis, our TUNEL provided a strong support to this notion. One plausible reason for this effect could be the modulation of WNT signaling pathway that had rendered DNA fragments.

While analysing the underlying molecular mechanisms of this quadruple-combinatorial treatment (TCNP), it is essential to clarify if the chosen drugs, both individually and as a combination could breach the BBB, a significant impediment for GB chemotherapy (Munoz Pinto et al. 2021). This is because, though several potent chemotherapeutic agents have been developed over the past decades, systemic administration of these drugs has not provided convincing benefits in prolonging patient outcome. Major part of this failure is attributed to the presence of the BBB, a highly vascularized interface that protects the neuronal microenvironment and also excludes chemotherapeutics from reaching the target site (Pandit et al. 2020). Thus, it is important to clarify if the chemotherapeutic agents could overcome the substantial barrier imposed by the BBB for effective GB therapy. This issue was addressed in our study, wherein, all the chosen drugs (both individually and as a combinatorial treatment) were detected either as parent ion or in their metabolic form in both the rat brain tissues and plasma samples (**Fig. 4, Fig. S2**). To note, so far, no report has described the simultaneous detection of all these drugs in GB bearing rat brain tissues. According to previous findings, TMZ was detected as its active metabolites, MTIC and AIC that had elicited its therapeutic effect against GB xenografts (Reyderman et al. 2004). However, this situation did not happen in our study; instead, TMZ was detected only in the form of its active metabolite AIC, while CHL was detected as desethylchloroquine. NAR and PGL, on the other hand, were detected in their parental state. This could be largely due to the instant entry of these drugs via the “short-circuit” routes which unchanged the detection pattern of these drugs. In this context, while this study provides preliminary evidence for the ability of the combinatorial treatment (TCNP) to cross the BBB and inhibit GB growth, our group is currently performing a more comprehensive study in GB-bearing drug-treated xenografts to quantitatively estimate the drug concentration and bioavailability for improvement of GB chemo-therapeutic efficacy.

Taken together, our *in vivo* findings had strongly emphasized that the quadruple-combinatorial treatment (TCNP) effectively hampered malignant glioma proliferation via modulation of WNT/β-catenin signalling cascade, paving way for intrinsic apoptosis in orthotopic GB-bearing xenografts. Nevertheless, there are certain limitations in the current study, that has to be looked upon in future, including quantitative estimation of drug levels in the treated GB xenografts and whether this quadruple-drug combination (TCNP) could also induce apoptosis extrinsically. Further, validation of the therapeutic efficacy elicited by this quadruple-combinatorial treatment (TCNP) in immunocompromised rodent models using TMZ-resistant cell lines will also be an interesting area of research to be thrived in mere future.

Nevertheless, lethality of GB depends not only on its proliferative ability, but also on several pathological features such as aberrant activation of signalling cascades and evasion of apoptosis. In the drug repurposing era for GB, the substitution of multi-drug cocktails for a single drug will provide superior benefit besides reducing toxicity. As a proof of concept, herein, we observed that the quadruple-combinatorial treatment (TCNP) was bestowed with an immense anti-glioma efficacy than the individual, dual or triple-drug treatments. Our *in vivo* evidence demonstrated that the quadruple-combinatorial treatment (TCNP) could synergistically inhibit GB growth and proliferation by altering WNT/β-catenin signalling pathway accompanied by apoptosis induction. These results strongly suggest that, the drugs CHL, NAR and PGL can be used in patient setting as a promising adjuvant regime to TMZ for GB treatment enhancement and patient’s well-being.

## ACKNOWLEDGEMENTS

This research work was supported by the Indian Council of Medical Research (ICMR) [Grant number: 5/13/21/2019/NCD-III]. The authors would like to thank Center for Stem Cell Research (A Unit of InStem, Bengaluru, India), Christian Medical College, Vellore and ESI-MS Instrumentation Facility, Pondicherry University, Puducherry for providing the instrumentation facility to carry out a part of this research. We would like to acknowledge Dr. Sowmya, Professor and Head, Department of Pathology, MGMCRI for her valuable inputs on pathological analysis. We also like to thank the Faculty of Mahatma Gandhi Medical Advanced Research Institute (MGMARI), Sri Balaji Vidyapeeth for giving us the basic infrastructure and instrumentation facilities.

