## Supplementary figures and images for "Dynamics of repurposing synthetic and natural compounds against WNT/β-Catenin signaling in glioma- An *in vivo* approach"

### Fig. S1

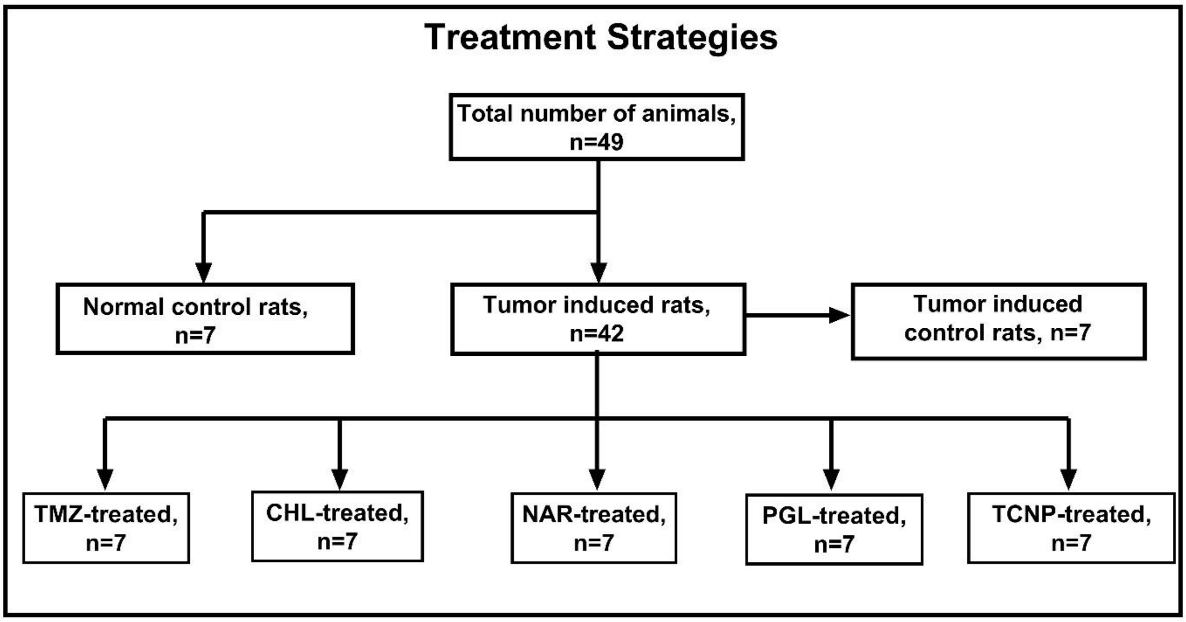


**Fig.S1** Treatment strategies employed for the study
