## Supplementary material for "Dynamics of repurposing synthetic and natural compounds against WNT/β-Catenin signaling in glioma- An *in vivo* approach": Fig. S2

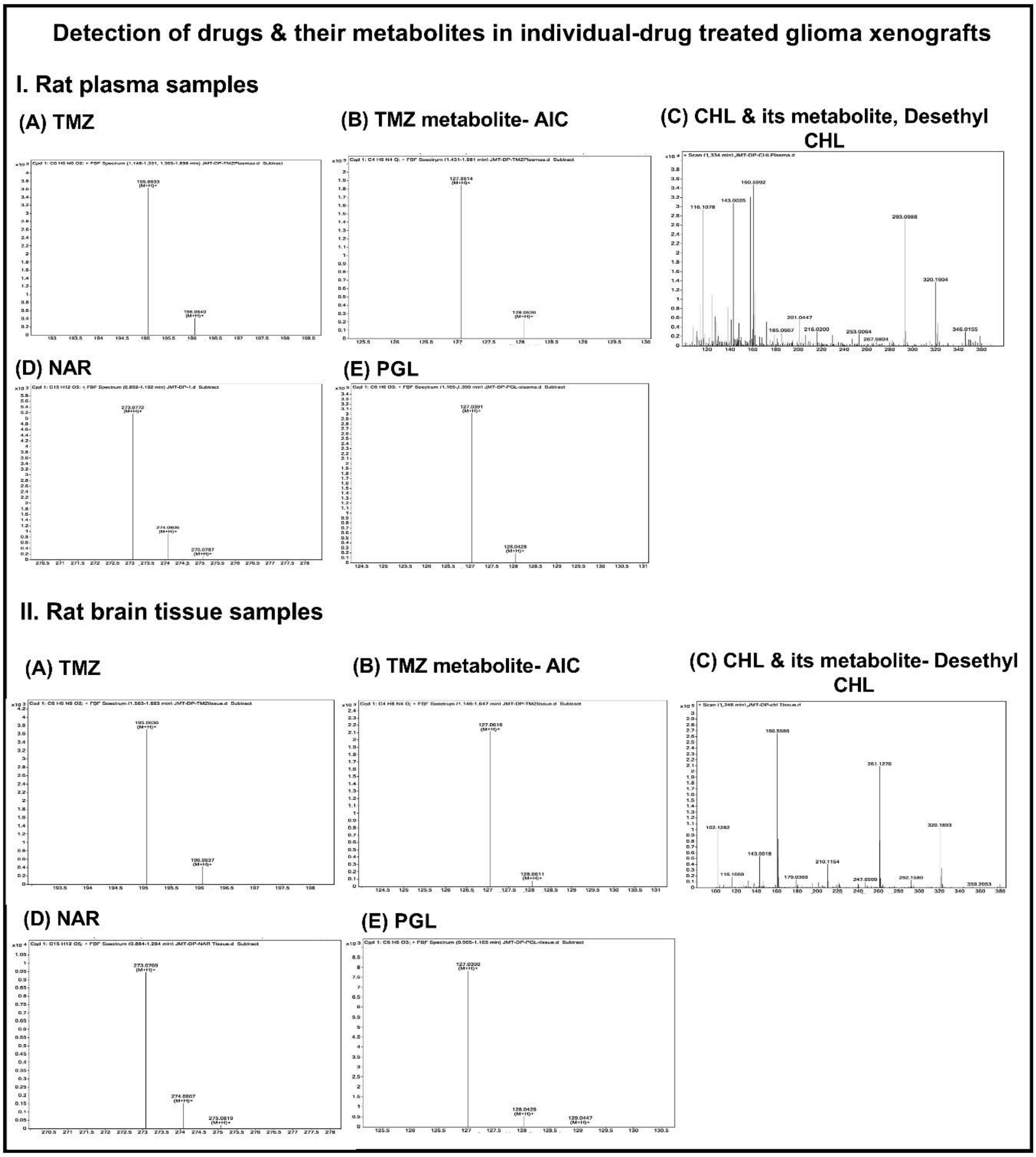


**Fig. S2 Ability of the individual drugs to breach the BBB and exert their therapeutic effect in GB-bearing xenografts**. Drug detection was assessed by ESI-Q-TOF MS in the GB-bearing -individual drug treated rat brain tissues and plasma samples
