## Supplementary material for "Dynamics of repurposing synthetic and natural compounds against WNT/β-Catenin signaling in glioma- An *in vivo* approach": Table S1

**Table S1** Human primer sequences employed for the study

| S. No | Primer | Forward Sequence (5'-3') | Reverse Sequence (5'-3') | Accession ID |
|---|---|---|---|---|
| **1** | EGFR | GGCACTTTTGAAGATCATTTTCTC | CTGTGTTGAGGGCAATGAG | NM_001346897.2 |
| **2** | β-catenin | TCTGAGGACAAGCCACAAGATTACA | TGGGCACCAATATCAAGTCCAA | NM_001330729.2 |
| **3** | c-myc | CCTGGTGCTCCATGAGGAGAC | CAGACTCTGACCTTTTGCCAGG | NM_002467.6 |
| **4** | c-jun | GCTCTGTTTCAGGATCTTGGGGTTAC | TTCTATGACGATGCCCTCAACGC | NM_002228.4 |
| **5** | ERK-2 | ACTCCTTTGAGCCGTTTGGA | AGTACATACTGCCGCAGGTC | Z_11694.1 |
| **6** | BAD | CGAGTGAGCAGGAAGACTCC | CACCAGGACTGGAAGACTCG | NM_032989.3 |
| **7** | cyt-c | TTCTAGGCACTGTCGGGGTA | AGCCTGAAGGCATGACGTTT | NM_018947.6 |
| **8** | Caspase-8 | CTGGTCTGAAGGCTGGTTGT | CAGGCTCAGGAACTTGAGGG | NM_001080124.2 |
| **9** | Caspase-3 | TGCTATTGTGAGGCGGTTGT | TCTGTTGCCACCTTTCGGTT | NM_032991.3 |
| **10** | BCL-2 | ATCGCCCTGTGGATGACTGAGTAC | AGAGACAGCCAGGAGAAATCAAAC | NM_000633.3 |
| **11** | β-actin | AGAGCTACGAGCTGCCTGAC | AGCACTGTGTTGGCGTACAG | NM_001101.5 |
